# Anatomical and functional organisation of the cholinergic nervous system in the tunicate *Botryllus schlosseri*

**DOI:** 10.64898/2026.09.10.750344

**Authors:** Chiara Anselmi, Leyla Yılmaz, Tom Levy, Katherine J. Ishizuka, Karla J. Palmeri, Irving L. Weissman, Ayelet Voskoboynik, Stuart Thompson

## Abstract

Acetylcholine (ACh) is an ancient, highly conserved neurotransmitter, yet the functional diversification of cholinergic pathways across early chordates remains incompletely understood. Here, we investigate the spatial and functional organization of the cholinergic system in the colonial tunicate *Botryllus schlosseri*, a chordate model that undergoes lifelong, cyclical neural regeneration. By integrating HCR-RNA FISH, in vivo pharmacology, extracellular brain electrophysiology, and quantitative reflex assays, we map the core molecular machinery of ACh synthesis (*ChAT)*, vesicular transport (*VAChT)*, *AChE*, and receptor signaling (*CHRNA7*, *CHRM3)*. Spatial expression analysis reveals a compartmentalized cholinergic network spanning central ganglia, peripheral sensory cells, and ciliated epithelia. Functionally, nicotinic receptors (nAChRs) mediate rapid mechanosensory burst firing in the brain, evoked siphon reflex contraction, and cilliary arrest, whereas muscarinic receptors (mAChRs), modulate baseline siphon motility and muscle tone. Pharmacological silencing of active siphon motor programs unmasks a slow, vascular-coupled rhythmic motility, while AChE inhibition induces paralysis indicating a critical role for regulated ACh breakdown and non-synaptic transmission. Together, these findings demonstrate that *B. schlosseri* possesses a sophisticated, dual-effector cholinergic system coordinating both muscular and ciliary networks, providing key insights into the evolutionary diversification of chordate neuromuscular control.

**SUMMARY STATEMENT:** By integrating spatial transcriptomics, electrophysiology, and in vivo pharmacology, this study reveals a compartmentalized cholinergic network in *Botryllus schlosseri* where nicotinic and muscarinic pathways differentially coordinate central burst firing, rapid siphon reflexes, basal muscle tone, and ciliary arrest.

## INTRODUCTION

### Acetylcholine: an ancient signaling molecule

Acetylcholine (ACh), a classical neurotransmitter, is an evolutionarily ancient signaling molecule found in bacteria, algae, plants, protists and animals (Wessler et al., 1999). In metazoans, ACh functions at neuronal synapses and neuromuscular junctions while also operating in diverse non-neuronal tissues to regulate homeostatic, ciliary and immune functions (Fujii et al., 2017; Tracey, 2002). At conventional cholinergic synapses, ACh is synthesized by choline acetyltransferase, packaged into synaptic vesicles by the vesicular acetylcholine transporter (VAChT) and rapidly released at specialized pre- and postsynaptic sites (Prado et al., 2013)(Fig.1A). In other cases, ACh functions as a paracrine signaling molecule in diverse neural and non-neuronal tissues (Proskocil et al., 2004) where it is released gradually from cells to act much like a hormone, a process referred to as non-neuronal transmission. Cholinergic signaling is mediated by two types of receptors, nicotinic ACh receptors (nAChRs) which are ligand-gated ion channels, and muscarinic ACh receptors (mAChRs), which are G-protein coupled receptors (Changeux, 2020; Hannan and Hall, 1993). The nAChR was the first neurotransmitter receptor to be isolated and fully described (Changeux, 2020; Tansey, 2006). Both have evolved a great deal of diversity in signal dynamics and molecular modes of action (Albuquerque et al., 2009; Changeux, 2020; Hannan and Hall, 1993; Pedersen et al., 2018).

**Figure 1:**
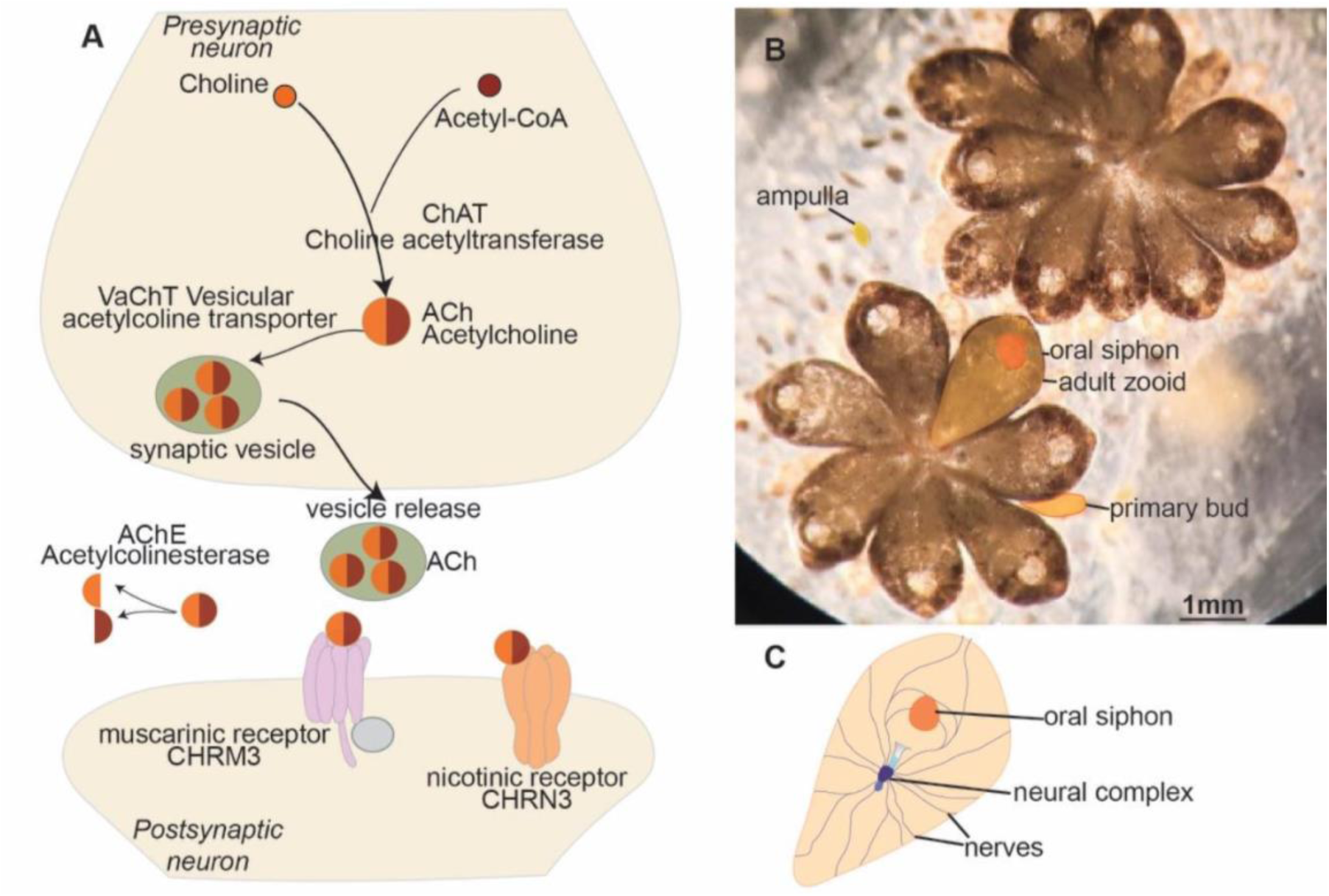
Overview of cholinergic signaling and the anatomical organization of *Botryllus schlosseri*. (A) Canonical cholinergic signaling pathway including the synthesis of acetylcholine by choline acetyltransferase (ChAT), vesicular packaging by VAChT, synaptic release, receptor binding to nicotinic (nAChR) and muscarinic (mAChR) acetylcholine receptors, and ACh degradation by acetylcholinesterase (AChE). (B) Bright-field image of a *B. schlosseri* colony composed of two systems. Each system contains multiple adult zooids with a visible open incurrent siphon and adjacent developing buds. (C) Dorsal view of an adult zooid showing the main nerves departing from the neural complex and projecting toward the oral siphon, musculature, and ciliated regions.

Despite the extensive characterization of cholinergic pathways in vertebrates and several invertebrate models, comparatively little is known about their organization and function in tunicates, the closest living relatives of vertebrates (Delsuc et al., 2006). Comparative studies in tunicates can therefore help distinguish ancestral features of chordate signaling systems from lineage-specific innovations that arose during vertebrate evolution (Anselmi et al., 2025a). Here, we investigate the cholinergic system of the colonial tunicate *Botryllus schlosseri* to examine how conserved molecular components of cholinergic signaling are spatially organized and contribute to neural and motor function.

### The tunicate *Botryllus schlosseri* as a model for cholinergic signaling and neuromuscular function

*B. schlosseri* is a colonial tunicate with a transparent body plan, compact genome, and a highly regular cycle of asexual development and regeneration (Levy et al., 2025; Manni et al., 2019; Vanni et al., 2022a; Voskoboynik and Weissman, 2015; Voskoboynik et al., 2013). Following settlement and metamorphosis, the larva gives rise to a sessile colony composed of genetically identical zooids, each approximately 2 mm long and containing a brain, peripheral nerves, heart, branchial basket, digestive system and gonads (Anselmi et al., 2025b; Kowarsky et al., 2021). Zooids are embedded within a common tunic and connected by an extracorporeal vascular network (Fig.1B).

A defining feature of *B. schlosseri* biology is its repeated cycle of blastogenesis. Under our culture conditions, each cycle lasts approximately seven days and includes three overlapping generations: adult zooids, primary buds and secondary buds (Fig.1B). At the end of each cycle, adult zooids undergo synchronous regression and are resorbed, while the primary buds become the new adult generation and secondary buds continue their development (Ballarin et al., 2010; Lauzon et al., 1993; Manni et al., 2019). This process, termed takeover, is repeated throughout the life of the colony, which can persist for years (Anselmi et al., 2022; Domen et al., 2026). Neural tissues are therefore repeatedly generated and remodeled across successive generations, making *B. schlosseri* particularly useful for studying nervous-system development and function in a recurrent developmental context (Anselmi et al., 2022; Anselmi et al., 2026; Bocci et al., 2024; Vanni et al., 2022b).

The adult cerebral ganglion contains approximately 900 neurons arranged as a peripheral cortical layer surrounding a central neuropil (Anselmi et al., 2022; Braun and Stach, 2019; Burighel et al., 2001; Bullock and Horridge, 1965). Adult brain development begins in the secondary bud and reaches maturity in the adult two weeks later, approximately three days after the siphons of the maturing zooids open and feeding begins. The number of neurons declines during the second half of the adult life cycle as takeover approaches (Anselmi et al., 2022; Anselmi et al., 2023). The brain is one component of a three part neural complex that includes the neural gland, and the dorsal organ and is located close to the oral siphon and ventral to the neural gland and dorsal organ and gives rise to nerve bundles that emerge from the anterior and posterior ends of the ganglion. The nerves lack myelination and typical glial cells but have a fibrous sheath. The anterior nerve roots split into smaller bundles that innervate the oral siphon and siphon tentacles. Histological staining for acetylcholinesterase (AChE) has long been used as a marker for nerves that link the *B. schlosseri* brain to muscles and internal organs (Arkett, 1987; Burighel et al., 2001; Mackie and Singla, 1983; Zaniolo et al., 2002). In some cases staining occurs along the entire length of the nerve in addition to regions known to be rich in synaptic terminals. An important conclusion from AChE histology is that there appears to be no neuron cell bodies in the periphery, the vasculature, the tunic or the branchial sac aside from sensory neurons. *B. schlosseri* zooids exhibit a great deal of motor activity involving internal muscle bands, body wall muscles, siphons, internal organs, heart and vasculature (Fig.1C). One conspicuous movement involves opening and closing of the incurrent siphon. These regular, but usually non-rhythmic, movements are part of filter feeding behavior driven by central motoneurons (Thompson et al., 2022).

Here, we combine HCR RNA-FISH, pharmacology, electrophysiology and behavioral assays to characterize the spatial organization and physiological roles of major cholinergic components in *B. schlosseri*. We examine the distribution of ChAT, VAChT, AChE, CHRNA7 and CHRM3 and test the contribution of nicotinic and muscarinic signaling, as well as acetylcholine breakdown, to spontaneous siphon activity, mechanically evoked reflexes and ciliary responses.

## MATERIALS AND METHODS

### Animal care and staging

Colonies of *B. schlosseri* (family Styelidae, order Stolidobranchia; Pallas, 1766), 0.5 - 2 years of age were obtained from the culture facility at the Hopkins Marine Station, Stanford Univ., Pacific Grove, CA., placed in a lab incubator and maintained in filtered sea water at 19^0^ C on a 11/13 dark: light cycle. The medium was exchanged and the colonies fed with rotifer culture every 4 days. Colonies were classified into stages A–D throughout the cycle.

Developmental stages were assigned according to (Watanabe, 1953). For recording, a colony adherent to a glass coverslip and bathed in filtered sea water was mounted on a temperature-controlled microscope stage (19^0^ C). Experiments were done between 11am and 4pm. Colonies returned to the incubator after recording could be maintained in healthy condition for several weeks permitting repeated experiments on the same colony over numerous cycles of blastogenesis. All experiments conformed to the animal care standards of Stanford University.

### Electrophysiology and pharmacology

Focal extracellular recordings were made using a polished glass micropipette (tip diameter 20-50 µm) filled with filtered sea water and fitted with a Ag:AgCl electrode. The recording pipette was pressed to the tunic directly over the brain or over an ampulla of the vascular system using gentle suction. Voltage was recorded differentially between the pipette and a Ag:AgCl electrode in the bath using AC-coupled preamplifiers (P55, Grass Instruments, Astro-Med, Inc., Warwick, RI). Signals were amplified by a factor of 1K or 10K and filtered between 30Hz and 1KHz (Frequency Devices, Ottawa, Il) before digitizing at 2KHz using a DATAQ Instruments DI-1000 AD-converter and WinDaq software (Dataq Instruments, Akron, OH). Digital records were analyzed with Igor Pro9 (WaveMetrics, Portland, OR). (+)-Tubocurarine chloride was from Sigma-Aldrich (93750) as was Acetylcholine chloride (<u>A6625</u>), Atropine sulphate (A0257) and Neostigmine bromide (N2001) were from Millipore-Sigma, Carbachol was from Calbiochem (212385). Pharmaceuticals were prepared in Micropure water (Barnsted) or in filtered natural sea water (0.45 µm Millipore Corp.). For bath application, the drug was added directly to the extracellular medium at concentrations between 30-200 µM. Prepared in this way, the drug enters the zooids with the incurrent sea water.

### Mechanical stimulation and drug delivery

Mechanical stimuli were delivered either as brief taps to the vibration-isolated recording table or as focal water-jet pulses. Water-jet stimuli were delivered through pulled glass micropipettes connected to a pneumatic microinjector (Picospritzer II, Parker Hannifin, Hollis, NH, USA).

### Video imaging

Colonies were viewed with a Wild stereo microscope fitted with a temperature-controlled stage and an AmScope (Irvine, CA) MD500 digital camera. Illumination was directed from above or below and images were collected using AmScope software at a resolution between 1280/1024 and 800/600 ppi and frame rate between 0.5 and 12.5 fps. Images stacks were analyzed using Fiji (ImageJ) and Igor Pro (WaveMetrics, Portland, OR). Changes in brightness, or the standard deviation of brightness, were measured within circular regions of interest (ROIs) positioned over the openings of oral siphons or over an ampulla of the vasculature. Contractions of zooids or ampullae resulted in an increase or decrease in reflected light intensity depending on the placement of the source and were quantified either as the change in average pixel intensity or the standard deviation of pixel intensity in the ROI as a function of time.

### Focal water jet stimulus to elicit siphon reflex

To quantify the sensitivity of the siphon reflex, we used a focal water-jet stimulation test adapted from (Anselmi et al., 2022). We performed a siphon stimulation test (SST) which involves mechanical stimulation of primary sensory cells located in the oral siphon wall with brief water jets delivered from glass needles (1 mm O.D., 0.78 mm I.D formed on a Sutter Instruments P-1000 micropipette puller and mounted on a manual micromanipulator (M3301R; World Precision Instruments). Tests were performed at constant temperature at the same time of the day. Water jets (impulses) were produced at approximately 1 minute intervals to allow the zooid to return to a relaxed condition and avoid habituation or sensitization. The pressure was gradually increased in 0.1 kPa steps starting from a minimum of 0.1 kPa, at which no behavioral response was observed.

Impulses were increased until the pressure was sufficient to cause oral siphon contraction at which point the pulse pressure was recorded. The expected reaction of zooids to the SST was the closure of the oral siphon. In controls the measurements were made after pressure injecting 5 µL of filtered natural sea water into the branchial basket and ampullae of a zooid. To study the effect of curare on the reflex, experiments were repeated after pressure injecting 5 µL of 30 mM D-Tubocurarine dissolved in distilled water into the branchial basket and ampullae of a zooid as before.

### Data and statistical analysis

For the focal water-jet stimulation assay, experimental subclones were treated as independent statistical units, whereas multiple zooids sampled within each subclone were considered within-subclone subsamples and were not treated as independent biological replicates. Individual zooids were not necessarily the same at the different time points; therefore, pre- and post-injection measurements were not paired at the level of individual zooids.

The original input data used for the analysis are provided in Supplementary Table 1. For each subclone and time point, the response threshold was summarized as the median pressure required to evoke oral siphon contraction among responding zooids. Zooids that failed to respond to the applied stimulus were classified as non-responsive and were not assigned a numerical threshold. Threshold comparisons between control and D-tubocurarine-treated subclones were performed using two-sided exact permutation tests on subclone-level median response thresholds. Because non-responsive zooids could not be included in numerical threshold analyses, responsiveness was analyzed separately. For each subclone and time point, the fraction of responding zooids was calculated as the number of responding zooids divided by the total number of zooids tested. Control and D-tubocurarine-treated subclones were compared using exact permutation tests on subclone-level responder fractions. Where time-specific follow-up comparisons were performed at both 15 and 60 min, P-values were adjusted for multiple testing using the Holm method. Individual zooid measurements are displayed in boxplots to illustrate the distribution of response thresholds, whereas statistical inference was performed using subclone-level summary values. Statistical significance was defined as *P*<0.05.

### Probe oligo-pool design for RNA HCR-FISH *in situ* hybridization

RNA probes for *BsACHE*, *BsCHRNA7*, *BsCHRM3*, *BsVAChT, BsCHAT* were obtained by insitu_probe_generator (ÖzpolatLab-GitHub-Kuehn, 2021, Opzolatlab-HCR,2021). In order to create the DNA oligo probe pairs specific to *B. schlosseri BsACHE*, *BsCHRN7*, *BsCHRM3*, *BsVAChT* messenger RNA *(mRNA)* (Voskoboynik et al., 2013), we used software (OzpolatLab_HCR, 2021) based on the probe design of HCR3.0 reported previously (Choi et al., 2014; Choi et al., 2018). B1 initiator (far red) is appended to each oligo in a pair. Exact oligo sequences are listed in the Supporting Information File. The sequences generated by the software were used to order a single batched DNA oligo pool (50 pmol DNAoPools Oligo Poll) from Integrated DNA Technologies. Details on the probe, amplifier and buffers are displayed in Supplementary Table 2. RNA HCR hairpins were ordered from Molecular Instruments (www.molecularinstruments.org). The sequence for HCR B1 amplifier is reported in (Choi et al., 2014).

### Tissue collection and HCR method

Tissue samples were fixed overnight in 4% paraformaldehyde at 4°C, washed three times in PBS, transferred to 100% methanol, and stored at −20°C until processing. Samples were rehydrated through a methanol/PBS series and HCR RNA-FISH was performed following the protocol of (Choi et al., 2014), using probe hybridization, probe wash and amplification buffers, and DNA HCR hairpin sets from Molecular Instruments. Samples were counterstained with DAPI before imaging.

### Imaging

HCR samples were mounted in a glass slide with DAPI, kept at 4℃ in the dark until imaging and imaged using a Zeiss LSM700 Confocal Microscope. Images were processed in FiJi (Schindelin et al., 2012).

## RESULTS

### Spatial organization of cholinergic pathway components revealed by HCR RNA-FISH

To investigate the spatial organization of the cholinergic system in *B. schlosseri*, we used HCR RNA-FISH to visualize the expression of five key genes, *ChAT*, *VAChT*, *AChE*, *CHRNA7*, and *CHRM3*, in the adult zooid body (Fig. 2). These genes collectively represent the core functional modules of the cholinergic pathway, from neurotransmitter synthesis and vesicular transport to receptor binding and transmitter clearance.

**Fig. 2:**
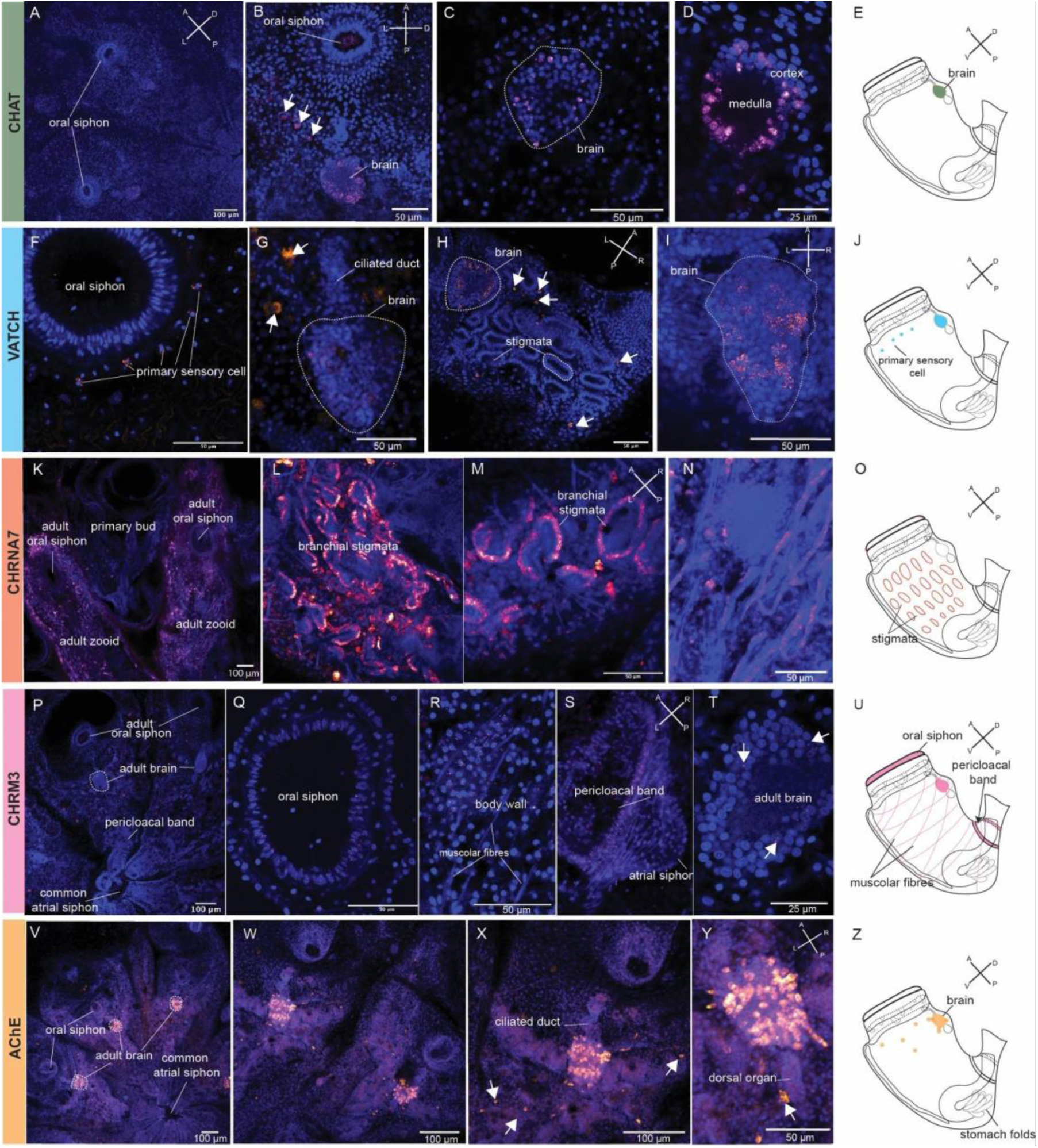
Spatial expression patterns of cholinergic pathway genes in *Botryllus schlosseri* zooids revealed by HCR RNA-FISH. Representative confocal images showing expression of five cholinergic pathway component gene-specific signals (magenta) together with DAPI nuclear counterstain (blue). Corresponding schematic illustrations summarize the principal expression domains observed for each gene in the adult zooid body plan. (A–E) ChAT (choline acetyltransferase): signal is detected in the cerebral ganglion (brain) and in adjacent dorsally positioned adjacent cell clusters (A–C; arrows), with higher-magnification views highlighting signal in the cortex surrounding the medullary region (D). A schematic overview of expression domains is shown in (E). (F–J) VAChT (vesicular acetylcholine transporter) signal is detected in primary sensory cells of the oral siphon epithelium (F) and in the adult brain (G). In the primary bud, additional puncta are visible in the dorsal epithelial (H, I; arrows). Panel (J) summarizes the main expression domains. (K–O) CHRNA7 (nicotinic acetylcholine receptor α7-like) signal is strongly enriched in the branchial stigmata (L, M), and associated peripheral epithelial regions. A detailed view is shown in (N). Panel (K) provides an anatomical overview including adult zooid and primary bud. A schematic representation is shown in (O). (P–U) CHRM3 (muscarinic acetylcholine receptor M3-like) signal is detected in muscle-associated territories, including the oral siphon (Q), body-wall muscle fibers (R), and pericloacal band and atrial siphon regions(S), with weak signal in the adult brain (T; arrows). An anatomical map of expression domains is provided in (U). (V–Z) AChE (acetylcholinesterase) signal is detected in the adult brain (V) and in additional peripheral territories, including body-wall cells (W; arrows), the ciliated duct (X), and the dorsal organ (Y). A schematic overview is shown in (Z). Scale bars are indicated in each panel.

ChAT, which encodes the enzyme that catalyzes acetylcholine synthesis, was strongly expressed in the cerebral ganglion and in adjacent cell clusters positioned dorsal to the ganglion whose identity remains unclear (Fig. 2A-E). Within the brain, the ChAT signal is not uniform but occurs in discrete domains, predominantly in the cortical layer (cortex) surrounding the central medullary region (medulla). At higher magnification HCR signal appeared mainly nuclear, with puncta concentrated within or closely overlapping the DAPI-defined nuclear area. Weak fluorescence was also detected in the developing brain of the primary bud, whereas no signal was observed in secondary buds.

The expression of VAChT, the vesicular acetylcholine transporter, was detected in the brain of adult zooids and buds (Fig. 2F–J). Peripherally, a clear VAChT signal was observed in primary sensory cells of the oral siphon epithelium (Fig. 2F). Additional VAChT-positive cells were detected in the primary bud (Fig. 2H). Notably, the arrow-marked signal is concentrated in the dorsal stigmatal region, consistent with primary sensory cells, although the identity of these cells cannot be assigned with certainty based on morphology alone. In contrast to ChAT, the VAChT HCR signal in brain cells appeared cytoplasmic rather than nuclear, with puncta distributed around the nuclei, compatible with cytoplasmatic/perinuclear distribution.

The nicotinic receptor alpha subunit *CHRNA7* showed a strikingly specific expression pattern, confined to the branchial stigmata, ciliated epithelial structures critical for filter feeding (Fig. 2K-O). In contrast to ChAT and VAChT, no CHRNA7 signal was detected in the brain, and labeling was instead confined to the peripheral epithelial compartment, with particularly robust expression around the neural complex in association with the stigmatal region. At higher magnification, CHRNA7 puncta appeared predominantly cytoplasmic rather than nuclear. CHRNA7 expression was not detected in the primary bud (Fig. 2K). This distribution is consistent with a peripheral cholinergic role in ciliated epithelial territories. CHRM3, a muscarinic acetylcholine receptor homolog, showed a complementary distribution biased toward muscle-associated territories (Fig. 2P–U). Robust expression was detected in the oral (Fig. 2Q) and atrial siphons and in the pericloacal band (Fig. 2S), where the signal was diffuse and closely associated with DAPI-positive nuclei. CHRM3 transcripts were also broadly present in body-wall muscle fibers, where labeling appeared widespread across the muscular layer (Fig. 2R). In addition to these peripheral sites, a weaker CHRM3 signal was observed in the cerebral ganglion (Fig. 2T). In this region, only a few cells displayed light staining, which appeared mainly cytoplasmic. No signal was detected in the buds.

AChE, the acetylcholine hydrolytic enzyme acetylcholinesterase, showed an anatomically organized expression pattern (Fig. 2V-Z). A strong HCR signal was detected in the cerebral ganglion, where labeling appeared predominantly nuclear, with puncta largely concentrated within the DAPI-defined nuclear area (Fig. 2X-Y). In contrast, along the nerves emerging from the brain, AChE staining was more cytoplasmic, with puncta distributed around nuclei and extending along the nerve profiles (Fig. 2Y). Beyond the neural complex, numerous AChE-positive cells were detected throughout the body wall. These cells (highlighted by arrows) occupy a position compatible with peripheral sensory elements, potentially including primary sensory cells, although their identity cannot be conclusively assigned based on morphology and location alone. Together, these data indicate that AChE is deployed both centrally and peripherally, with distinct subcellular localization in the brain versus peripheral nerves and a prominent population that may participate in sensory or neuroepithelial cholinergic regulation.

Altogether, the HCR RNA-FISH data reveal a distributed and compartmentalized cholinergic system in *B. schlosseri*, spanning both central and peripheral domains. The distinct expression patterns of biosynthetic, receptor, and degradative components indicate spatial differentiation within the cholinergic pathway in this basal chordate.

### D-tubocurarine suppresses spontaneous siphon closure and alters siphon muscle tone

To test whether nicotinic ACh receptors contribute to ongoing siphon behavior, we examined the effects of D-tubocurarine (curare). Curare is a competitive inhibitor of acetylcholine binding to nAChR that has little effect on mAChR. We examined its effect on neural activity in the *B. schlosseri* brain and on siphon behavior by microinjecting curare into vascular ampullae by micropipette. On injection, it enters the open circulation of the zooids to reach the brain, siphon muscles, and sensory neurons associated with the siphon and coronal organs. We found that curare prevents the spontaneous opening and closing of the oral siphons that are a key part of feeding behavior (Fig. 3). In addition, the apertures of the siphons appeared relaxed and expanded compared to control conditions. In contrast, curare did not affect heart beat or the steady rhythmic contraction of vascular ampullae.

**Figure 3:**
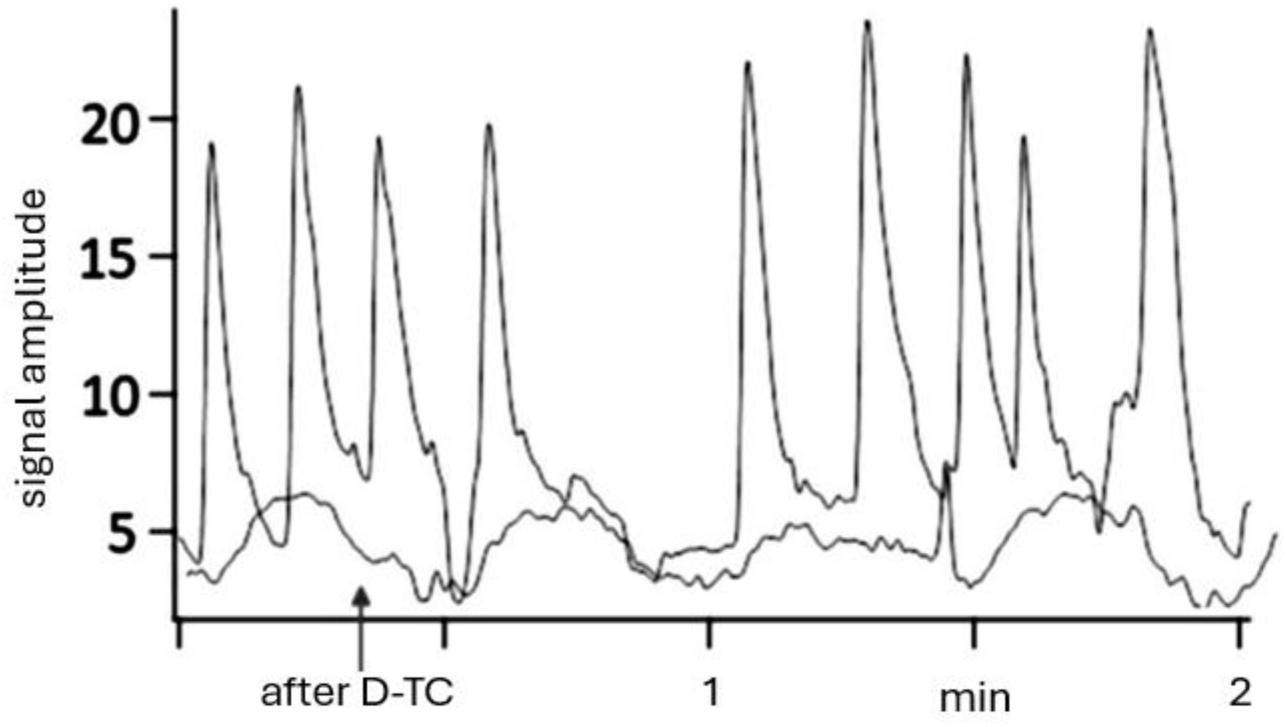
Spontaneous siphon closure is blocked by curare exposing a slow rhythm. Video recordings of siphon motility were made after assigning ROIs to the apertures of incurrent siphons. The upper trace shows control siphon closures in a single siphon. In this example, closure results in an increase in reflected light intensity. The lower trace is a recording from the same ROI 15 min after microinjecting ∼ 30 µM curare into three adjacent vascular ampullae. Shortly after injection the siphon muscles begin to relax and spontaneous siphon closures cease. At the same time, a smaller, slower movement is exposed (label; after D-TC) that occurs with a period of about 30 sec, approximately the same as the ampullae rhythm. The two records are plotted on the same vertical scale. (Zooid stage A3.)

The effect of curare is slowly reversible. We observed that after curare the entire zooid appeared to relax and to exhibit slow, low amplitude movements with a period of about thirty seconds, a period similar to the regular rhythm of vasculature contractions. The slow movements could be recorded with video imaging at many places on the body, including at the oral siphon (lower trace in Fig. 3). The source of the slow rhythm is not clear, but we suggest that vascular contractions might pull on the now relaxed smooth muscle of the body via the tunic to passively generate the movement (De Santo and Dudley, 1969; Thompson et al., 2022). Because curare was delivered through the vasculature, these experiments do not distinguish between central and peripheral sites of action. Thus, the observed behavioral effects may result from inhibition of nicotinic signaling in the brain, at neuromuscular junctions, or both.

### Curare alters spontaneous neural activity recorded from the brain

Focal extracellular recording was used to measure neural activity in the brain under control conditions and after microinjecting ∼30 µM curare into the vascular system (Fig. 4). Curare is judged to have reached an effective blocking concentration at nAChRs once spontaneous siphon closures had ceased and the siphons no longer respond to mechanical stimulation.

**Figure 4:**
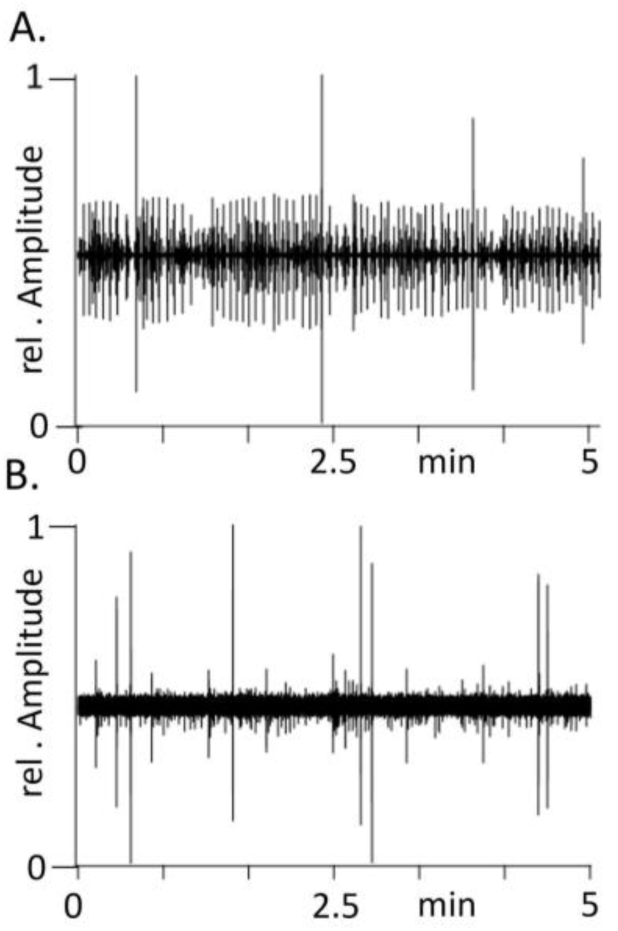
Spontaneous neural activity in the *B. schlosseri* brain before and after introducing curare. Neural activity was recorded with a focal extracellular electrode appressed to the tunic over the brain by gentle suction. (A) Action potential activity in the brain under control conditions. (B) The recording was repeated at the same position 30 minutes after the microinjection of curare into vascular ampullae to an estimated final concentration of ∼30 µM. With this method, curare has direct access to the brain via the open circulation of the zooid. (Zooid stage A3).

Under control conditions, extracellular recording reveals complicated patterns of spontaneous electrical activity composed of action potentials of varying amplitude, indicating contributions from a number of different types of neurons. Curare did not eliminate all neural activity in the brain. Instead, it caused a dramatic change in firing pattern, decreasing the overall frequency of action potentials, and blocking of action potentials in certain classes of neurons, as judged by the elimination of spikes of intermediate amplitude, while sparing others. We propose that the activity that is blocked by curare originates in neurons that normally participate in nicotinic neural circuits.

### Nicotinic signaling is required for the mechanically evoked siphon reflex

We next tested the role of ACh in the siphon reflex evoked by mechanical stimulation*. B. schlosseri* responds to a brief mechanical stimulus, for example a 250 msec mechanical tap to the metal isolation table, with reflex closure of the incurrent siphons that begins with a delay and is preceded by a characteristic burst of action potentials in the brain (Fig. 5A, (Thompson et al., 2022). We examined the role of nAChR in the siphon reflex by injecting ∼30 µM curare into vascular ampullae. Following curare injection, both the behavioral response (Fig. 5B) and the associated action potential burst are strongly reduced or abolished (Fig. 5C-D). We do not know if this is due to a block of the afferent sensory input to motor neurons in the brain or an inhibitory action at another place in the reflex pathway. We note, however, that the neural response to the mechanical stimulus in control conditions is quite prolonged, lasting more than twenty seconds in Fig.5C, and involves multiple classes of neurons, based on action potential amplitudes. This suggests that the sensory input from a brief stimulus spread widely and may activate multiple neural circuits. In some experiments, siphon contraction was completely blocked by curare, but a weaker neural response remained that might be due either to incomplete inhibition of nAChR, or possibly from sensory activation of unrelated neural pathways.

**Figure 5:**
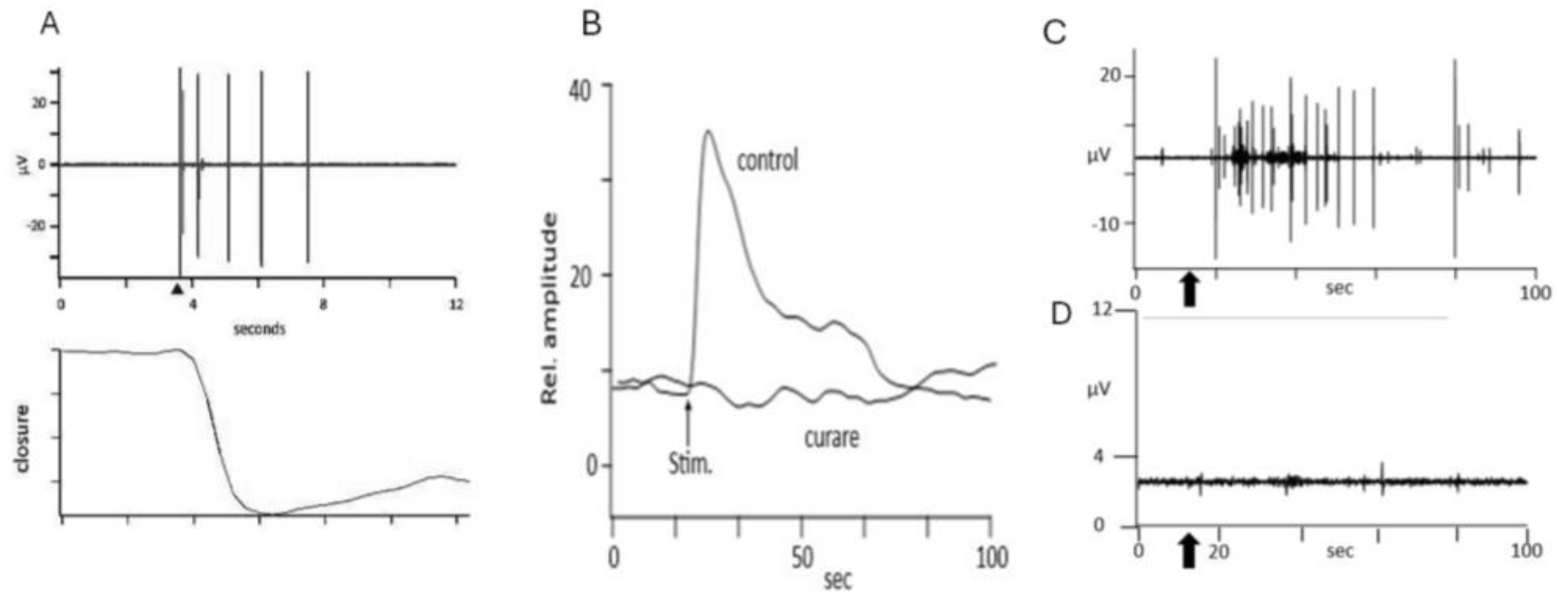
The siphon reflex in response to mechanical stimulation is blocked by curare. (A) The siphon reflex in control conditions (zooid stage A4). The upper trace shows neural activity recorded from the brain in response to a sharp tap applied to the table top (arrow) by a metal solenoid. The lower trace shows a video record of the closure of the incurrent siphon recorded at the same time. Closure begins with a delay of about 1 sec. In this example, siphon closure causes a decrease in reflected light intensity. (B) Reflex contraction of the siphon in the control and 15 minutes after injecting curare into adjacent vascular ampullae to an approximate final concentration of ∼30 µM. Curare eliminated the reflex response (zooid stage A3). (C) Electrical activity in the brain in a different preparation (zooid stage A3). The upper trace shows the prolonged neural response to a mechanical tap (arrow). (D) The lower trace shows a repeat of the experiment 20 min after microinjecting curare. Curare eliminated the action potential burst in central neurons. Zooid stage A3.

### Focal water-jet stimulation confirms impairment of the siphon reflex after curare treatment

To define the sensory pathway involved in siphon closure more precisely, we used a focal water-jet stimulation assay directed at the oral siphon epithelium. Unlike the table-tap stimulus, this assay provides a more localized mechanical stimulation of the oral siphon region. Under control conditions, brief water pulses triggered contraction of the oral siphon once a threshold pressure was reached. The measurement was repeated after injecting curare into both the zooid body cavity and into adjacent ampullae (Fig. 6A). Fifteen minutes after D-tubocurarine injection, response thresholds were significantly higher than in control subclones (*P*=0.0357; Fig. 6B). The fraction of responding zooids was also markedly reduced, with partial recovery by 60 min (Fig. S1), although time-specific differences in responder fraction were not significant after Holm correction. At 60 min, response thresholds remained significantly elevated relative to controls (*P*=0.0083; Fig. 6C). These results independently support the conclusion that nicotinic cholinergic signaling is required for normal sensitivity and motor execution of the siphon reflex.

**Fig 6:**
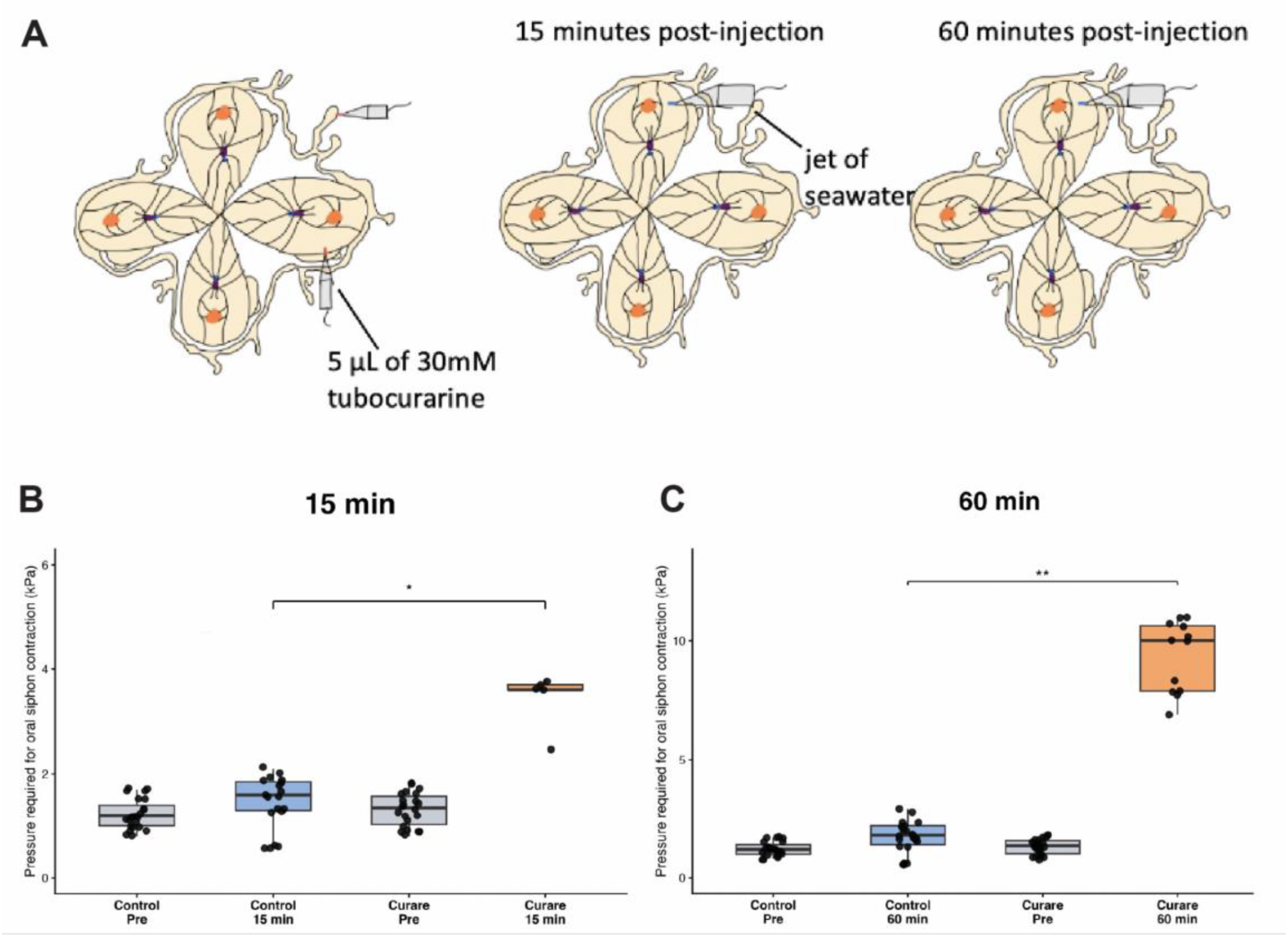
Focal water-jet stimulation assay shows increased response threshold after D-tubocurarine treatment. (A) Experimental design for focal water-jet stimulation. Stage A subclones were first tested before injection and then injected either with 5 µL of filtered natural seawater (control) or with 5 µL of 30 mM D-tubocurarine. Oral siphon responses to focal seawater stimulation were assessed 15 and 60 min after injection. (B) Response threshold 15 min after injection. Individual zooid measurements are shown for visualization; boxplots represent the distribution of pressure values required to elicit oral siphon contraction in responding zooids. D-tubocurarine- treated subclones showed a higher response threshold than control subclones at 15 min (*P* = 0.0357). (C) Response threshold 60 min after injection. D-tubocurarine-treated subclones showed a markedly elevated threshold compared with controls (P = 0.0083). Only responding zooids are included in the threshold boxplots. Statistical comparisons were performed at the subclone/experimental-series level using two-sided exact permutation tests on subclone-level median response thresholds, with multiple zooid measurements within each subclone treated as subsamples. *P < 0.05; ** P < 0.01;

### The mAChR antagonist atropine reduces spontaneous siphon motility but does not block the neural response to mechanical stimulation

To examine the contribution of mAChR to motor behavior we applied atropine to the external medium and monitored spontaneous siphon motility and the mechanically evoked refex clousure. Atropine is known to be a competitive antagonist of acetylcholine at all five subtypes of muscarinic Ach receptors (M1 to M5) found in the *Botryllus* genome. Atropine sulphate (50-200 µM) was added in the external medium. It enters the zooids with the incurrent water flow to access the brain, internal muscles and organs, and sensory receptors. Applied in this way, 50 µM atropine reduced the frequency and amplitude of spontaneous siphon closures (Fig. 7A), and exposed a slow, low amplitude movement similar to what is seen with curare (Fig.3), but it did not inhibit all siphon motility. We observed, however, that atropine had little effect on reflex siphon contractions in response to mechanical stimulation and the same result was obtained when the atropine concentration was increased to 200µM. We used mechanical taps to stimulate reflex siphon closures while recording electrical activity from the brain. The neural response under control conditions is illustrated in Fig. 7B which shows that a single tap (arrow) results in a burst of action potentials that began with a significant delay. The experiment was repeated 45min after adding 50 µM atropine sulphate to the external medium and it was found that the mechanical stimulus continues to activate a neural response, again with a long delay (Fig. 7C).

**Figure 7:**
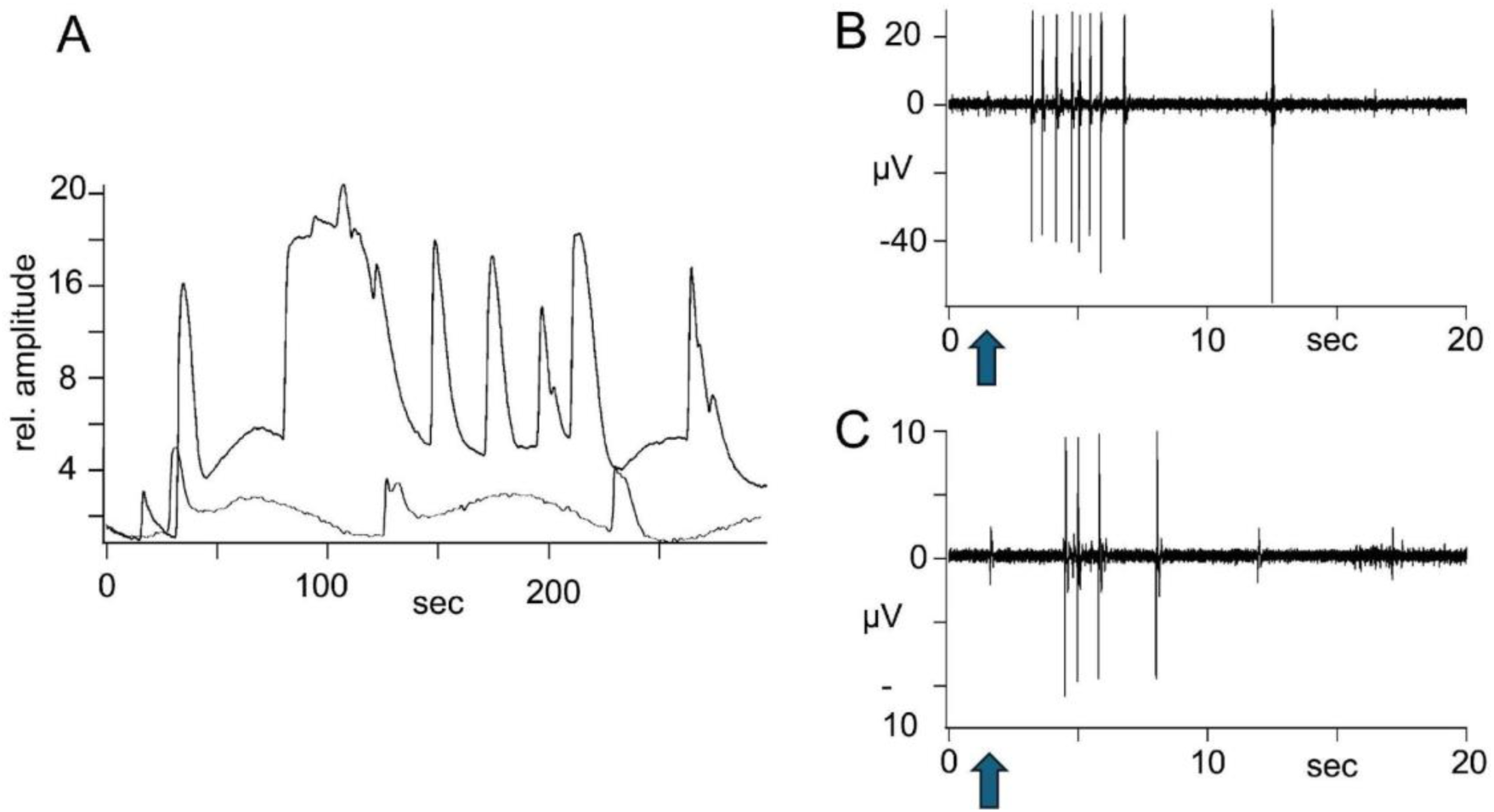
Atropine modifies ongoing siphon closures associated with feeding behavior but does not eliminate the siphon reflex . (A) The upper trace shows video recording of spontaneous siphon closures in control conditions (closures shown as upward deflections). Atropine sulphate (50 µM) was then added to the external medium (lower trace). After 45 min, video recording was repeated and showed that regular siphon contractions were greatly reduced or eliminated. In addition, atropine exposed a much smaller, slower movement similar to what was seen in the presence of curare. (B) Electrical recording from the brain in a different preparation under control conditions showing the burst of action potentials in response to a brief mechanical tap (arrow). **C:** The mechanical stimulus (arrow) was repeated 60 min after adding 50 µM atropine to the external medium. The stimulus continued to elicit a burst of action potentials, again after a substantial delay. (zooid stage A3-A4)

### Inhibition of AchE with neostigmine suggests a role for non-neuronal cholinergic signaling

We next examined the effect of neostigmine, a reversible inhibitor of acetylcholinesterase (AChE) that competes for binding sites on the enzyme, on siphon behaviors. AChE is widely expressed in *B. schlosseri,* not only at synaptic sites but along the length of nerves and in the vicinity of non-neuronal cells, especially ciliated cells in the stomach and stigmata and in the vicinity of the siphon tentacles (Fig.2). In clinical practice, neostigmine and similar AChE inhibitors are used to strengthen the response at ACh dependent neuronal synapses and neuromuscular junctions by lengthening the exposure to ACh. In our experiments on siphon smooth muscle, however, neostigmine had the opposite effect. Instead of enhancing reflex contraction, neostigmine, (33 µM) applied in the external medium, caused muscles of the zooid body and the siphons to relax and largely blocked the siphon reflex in response to mechanical stimulation (Fig.8). This resembles a well-known clinical effect of high neostigmine concentration termed cholinergic crisis, a state of muscle paralysis due to excess accumulation of ACh at neuromuscular junctions leading either to depolarization block (Bianchi et al., 2012) or to receptor desensitization. We suspect that the cholinergic crisis may explain our results, although we could not directly test this idea. The observation that neostigmine causes paralysis suggests that under normal conditions gradual but continuous non-synaptic release of ACh, that is normally balanced by AChE hydrolysis, may play an important role in establishing muscle tone and that inhibition of AChE with neostigmine may allow ACh to accumulate to excess, paradoxically lowering muscle tone.

**Figure 8:**
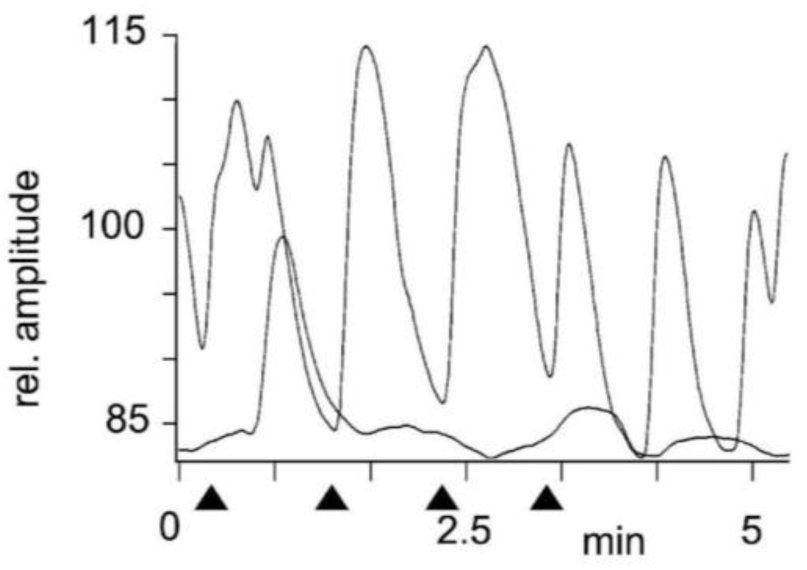
Effect of neostigmine on spontaneous and evoked siphon activity. Video recordings of siphon closure under control conditions (larger trace) and 13 min after adding 33µM neostigmine to the bath (smaller trace). The two records are plotted on the same vertical scale. Neostigmine greatly decreased siphon contractions. It also exposed a slow, low amplitude contraction rhythm that resembles what is seen after blocking nAChR or mAChR. Reflex closure of the siphon in response to mechanical stimulation is also prevented by neostigmine. Four mechanical taps were applied before (larger trace) and again after adding 33µM neostigmine to the external medium. The timing of taps is shown by arrowheads, and the stimuli were applied with the same timing in control and after applying neostigmine. In the control, each tap caused reflex closure of the siphon after a brief delay. There was no response to any of the four taps after neostigmine.

### Cholinergic signaling produces induces ciliary arrest in feeding-related tissues

*Botryllus* zooids are filter feeders, capturing small particles from the water that enters through the incurrent siphon. Cells in the branchial basket bear motile cilia that maintain a regular beat to move food particles through the pharyngeal gill slits (stigmata). The cilia in these organs experience ciliary arrest and stop beating for a time as part of the reflex response to mechanical stimulation (Mackie et al., 2006). We used video microscopy to confirm that ciliary beating in the gut and tentacles is interrupted by vibrational stimuli, and we found that direct application of 10µM acetylcholine or carbachol, a cholinergic agonist, to ciliated cells of the digestive system evokes ciliary arrest. The area adjacent to these cells stains strongly for AChE (Fig.2). These findings are consistent with those of (Arkett, 1987) who used intracellular recording to demonstrate that direct depolarization of specific neurons in the brain of a related tunicate (*Corella inflata*) causes ciliary arrest in the branchial sac. Also, (Jokura et al., 2020) reported that an alpha 7-related subunit of nAChR mediates ciliary arrest in the gut of the tunicate *Ciona..* It is likely that in the *B. schlosseri* nervous system, the muscular siphon reflex and ciliary arrest represent two parallel cholinergic pathways activated by mechanosensory input.

## DISCUSSION

### Cholinergic pathways in *Botryllus schlosseri*: integration of anatomical, molecular and functional evidence

Our results extend earlier morphological studies showing peripheral nerve fibers arising from the cerebral ganglion and innervating major visceral, muscular, and ciliated territories, with no clear peripheral neuronal somata (Burighel et al., 2001; Zaniolo et al., 2002). These projections develop early in blastogenesis and are extensively remodeled later in the cycle. The sequential emergence of cholinergic components across blastogenic generations, beginning with the early expression of biosynthetic and transport machinery (*ChAT* and *VAChT*) in the developing primary bud prior to the full deployment of mature effector receptors (*CHRNA7* and *CHRM3*) in functional adults, indicates that cholinergic circuitry undergoes progressive synaptogenesis and maturation that coincide with siphon opening and the onset of active filter feeding.

Our HCR RNA-FISH data are broadly consistent with this anatomical framework, while providing gene-specific information on the distribution of cholinergic components. ChAT was strongly expressed in the cerebral ganglion and major nerve-associated territories, consistent with its role in ACh biosynthesis in cholinergic neurons (Oda, 1999; Pereira et al., 2015; Takamura et al., 2010). VAChT was also detected in the brain, supporting the presence of vesicular cholinergic transmission machinery (Prado et al., 2013).

Within the cerebral ganglion, the prominent localization of *ChAT* and *AChE* puncta within or overlapping DAPI-positive nuclear boundaries likely reflects active foci of nascent pre-mRNA transcription within the compact nuclei of ascidian neurons, contrasting with the predominantly cytoplasmic distribution of mature receptor transcripts observed in peripheral tissues.

Notably, ChAT- and VAChT-associated signals were also detected outside the central ganglion, including regions surrounding the brain and peripheral territories. Previous anatomical studies in *B. schlosseri* found no clear peripheral motor neuronal somata (Burighel et al., 2001; Zaniolo et al., 2002) whereas in *C. intestinalis*, peripheral neural cells can occur in small miniganglia along major nerves (Dahlberg et al., 2009). Our data therefore, raise the possibility that peripheral cholinergic cell populations may also occur in *Botryllus*, although higher-resolution analyses will be required to determine whether the observed signals correspond to neuronal somata, axonal compartments, or other cholinergic cell types.

AChE showed a broad distribution, including muscle- and cilia-associated regions such as the oral siphon, consistent with cholinergic neuromuscular signaling in ascidians (Brown and Piscopo, 2011; Takamura et al., 2002). Receptor expression was also spatially differentiated. CHRNA7 was enriched in branchial stigmata and associated epithelial territories, whereas the CHRM3-like receptor was enriched in muscle-associated and visceral regions. In vertebrates, α7-containing nicotinic receptors participate in neuronal signaling and plasticity (Albuquerque et al., 2009) while M3-type muscarinic receptors regulate smooth muscle contraction and glandular secretion (Eglen, 2006). Tunicate muscarinic receptors are phylogenetically close to ancestral lineages that gave rise to vertebrate mAChR subfamilies, which diversified after the tunicate–vertebrate split (Pedersen et al., 2018).

Together, these findings reveal a spatially compartmentalized cholinergic system in *B. schlosseri*, in which central, peripheral, epithelial, and muscle-associated domains are molecularly distinct and positioned to support different aspects of neural and effector function.

### Cholinergic regulation of siphon motility and reflex behavior

*B. schlosseri* is a filter feeder in which ciliary beating and siphon movements regulate water flow. D-tubocurarine reduced spontaneous siphon closures and mechanically evoked responses, while altering spontaneous neural activity and abolishing the evoked neural burst, supporting a major role for nicotinic signaling in both ongoing and stimulus-evoked motor behavior. This is consistent with previous studies showing that siphonal mechanosensory cells contribute to mechanically evoked responses in ascidians (Anselmi et al., 2022; Anselmi et al., 2024; Mackie and Wyeth, 2000; Mackie et al., 2006; Manni et al., 2018) and that spontaneous and evoked neural activity accompanies siphon movements in *Botryllus* (Thompson et al., 2022).

In the focal water-jet assay, D-tubocurarine-treated zooids required stronger stimulation to contract the oral siphon, and many failed to respond within the tested range. Because the drug was delivered through the vasculature and body fluids, these experiments cannot distinguish effects on peripheral sensory input, central circuits, neuromuscular targets, or multiple sites. They nevertheless establish a requirement for nicotinic signaling in normal siphon reflex function. D-tubocurarine also altered, but did not abolish, spontaneous cerebral ganglion activity, suggesting disruption of specific circuit components rather than global suppression of neural function. This is consistent with larval *Ciona*, where cholinergic motor neurons express ChAT and VAChT and muscle nAChRs are required for normal motor control and swimming behavior (Nishino et al., 2011; Takamura et al., 2002).

Mechanical stimulation also evoked activity that far outlasted the stimulus itself, suggesting recruitment of sustained sensory or central circuitry. Such activity could coordinate multiple mechanically sensitive outputs, including siphon contraction and ciliary arrest, although the underlying circuit organization remains unresolved.

By contrast, atropine reduced the amplitude and frequency of spontaneous siphon closures but did not abolish the mechanically evoked neural response. Muscarinic signaling may therefore contribute more strongly to baseline motor state and muscle tone than to the rapid evoked neural response. This interpretation is consistent with the predominantly peripheral and muscle-associated distribution of the CHRM3-like receptor. Muscarinic regulation of contractile tissues is widespread in vertebrates (Eglen, 2006) and homologous muscarinic receptor systems occur across diverse invertebrates (Hannan and Hall, 1993). Taken together, these pharmacological and electrophysiological distinctions point toward a dual-component neuromuscular model in *Botryllus*: a fast, phasic nicotinic pathway mediating rapid mechanosensory signal transmission, central burst dynamics, and swift reflex contractions, operating alongside a slower, modulatory muscarinic (*CHRM3*) pathway that fine-tunes resting baseline muscle tone and visceral motility.

AChE inhibition with neostigmine also reduced spontaneous and mechanically evoked siphon activity. Because AChE inhibition prolongs ACh availability (P. Taylor, 2011), this phenotype suggests that coordinated siphon behavior requires tightly regulated ACh clearance rather than sustained cholinergic stimulation.

A slow residual siphon movement was observed following D-tubocurarine, atropine, and neostigmine treatment. Its periodicity resembled vascular ampullar contractions described in *Botryllus* (Thompson et al., 2022). One possibility is that this movement normally remains masked by stronger zooid motor activity and becomes apparent when cholinergic-dependent muscle tone is reduced. Although indirect, this interpretation suggests a mechanical contribution of the vascular rhythm to residual zooid movement rather than a component of the normal cholinergic siphon motor program. This biomechanical unmasking highlights the functional interplay between active muscular tone and the colonial circulatory system, wherein the tonic contractile state of zooid smooth muscles normally shields body apertures from the passive rhythmic mechanical forces exerted by continuous vascular ampullar contractions (Thompson et al., 2022).

Together, these pharmacological effects suggest distinct cholinergic contributions to behavior in *B. schlosseri*: nicotinic signaling is particularly important for rapid sensorimotor output, muscarinic signaling contributes to ongoing motor state and muscle tone, and AChE-dependent ACh clearance supports coordinated responses. More broadly, the behavioral effects observed in *B. schlosseri* parallel pharmacological responses described in both vertebrates and other invertebrates, supporting the evolutionary conservation of cholinergic sensitivity across animal systems.

### Peripheral cholinergic regulation, ciliary control, and evolutionary implications

AChE histochemistry and HCR RNA-FISH indicate cholinergic-associated territories extending beyond the cerebral ganglion. Earlier ultrastructural studies also described axonal varicosities without clear conventional synaptic specializations (Arkett, 1987; Burighel et al., 1998; Burighel et al., 2001). Together, these observations raise the possibility that cholinergic signaling in *Botryllus* may include non-synaptic or paracrine components, although this remains to be demonstrated directly.

Non-neuronal cholinergic signaling is widespread in vertebrates, and organic cation transporters (OCTs) can mediate ACh transport in some systems (Kawashima and Fujii, 2000; Wessler et al., 2001; Wessler et al., 2003). The presence of OCT homologs in *B. schlosseri*, together with the effects of cholinergic antagonists on muscle tone, therefore raises the possibility of tonic, non-synaptic peripheral ACh signaling.

In such a system, non-synaptic release via axonal varicosities or OCT-mediated transport could operate through paracrine volume transmission, wherein a homeostatic equilibrium between continuous diffuse ACh release and rapid AChE hydrolysis establishes a basal threshold of tissue excitability across the colony.

Cholinergic regulation also extends to ciliated effectors. CHRNA7 was enriched in branchial stigmata and epithelial territories, and ACh or carbachol induced ciliary arrest. This is consistent with α7-related nicotinic control of ciliary arrest in *Ciona* (Jokura et al., 2020) and earlier observations linking neural activity to inhibition of ciliary beating in ascidians (Arkett, 1987). Cholinergic signaling in *B. schlosseri* may therefore coordinate both muscular and ciliated effectors involved in feeding and defensive responses.

The recurrent blastogenic renewal of *B. schlosseri* provides a distinctive context in which to examine how this conserved signaling system is repeatedly established and remodeled. Future work should resolve the identity of peripheral cholinergic marker-positive cells, define the anatomical sites of drug action, and directly test whether non-synaptic ACh signaling contributes to peripheral regulation.

**Table 1.**
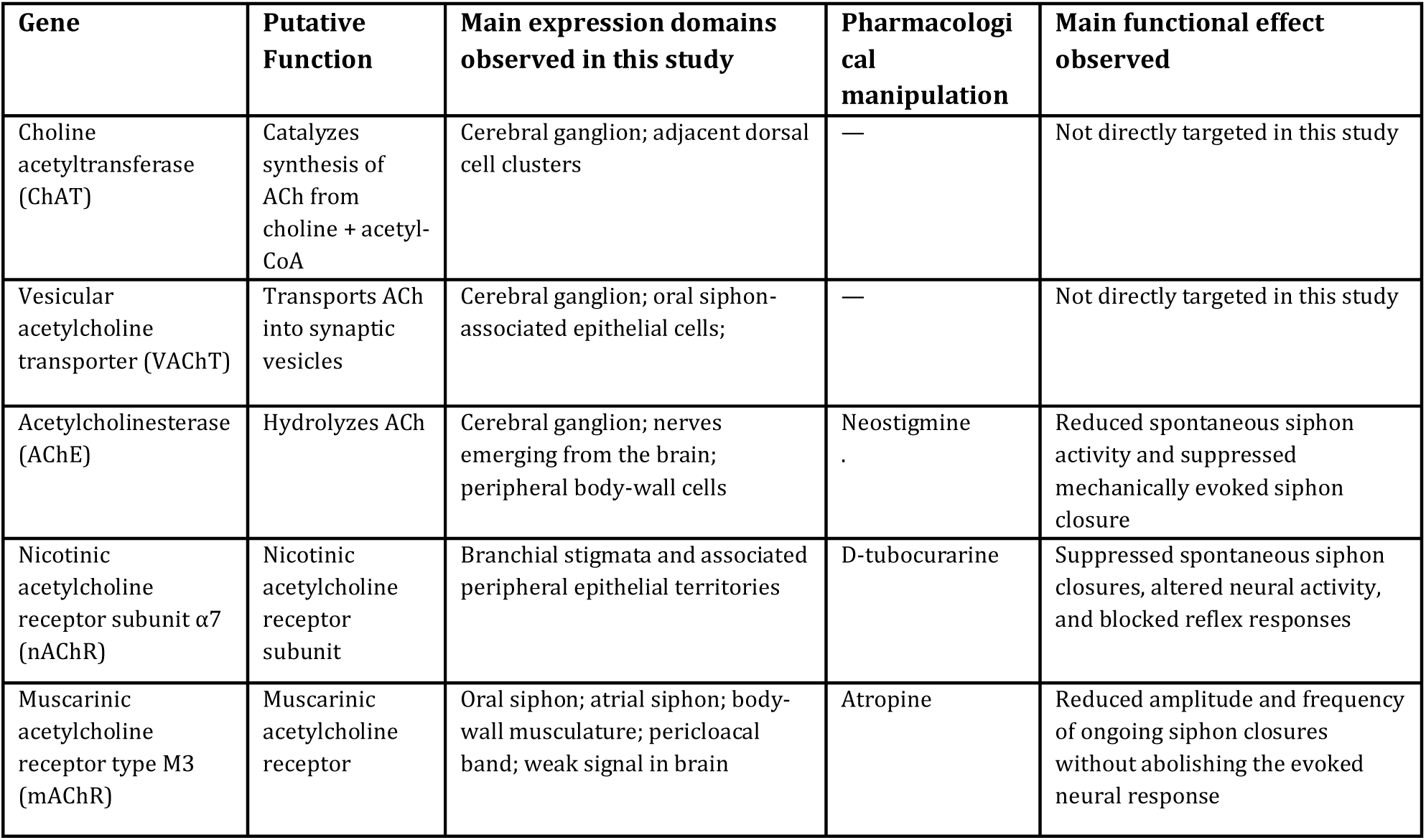
Summary of cholinergic agents and genes used to study the cholinergic system in *B. schlosseri*. The table summarizes the cholinergic genes examined in this study, their inferred functions, the main expression domains observed by HCR RNA-FISH, the pharmacological agents used to perturb cholinergic signaling, and the principal behavioral or physiological effects observed in vivo. Expression domains and functional effects are limited to the data presented in this study.

## AUTHOR CONTRIBUTION

C.A., A.V., and S.T. conceived the study, and designed the overall project. C.A. and S.T. interpreted the results, and wrote the first draft of the manuscript. C.A., L.Y., and T.L. performed the HCR RNA-FISH experiments. C.A. and L.Y. performed the siphon stimulation test, associated behavioral analyses, drug injection and statistical analysis. S.T. performed the electrophysiological recordings, video imaging, and pharmacological experiments. K.J.I. and K.J.P. contributed to animal care, colony maintenance, and experimental support. ILW and A.V. provided essential laboratory resources, and facilities A.V. provided genomic models. All authors read and approved the final manuscript.

## SUPPLEMENTARY MATERIAL

**Supplementary Fig. S1.**
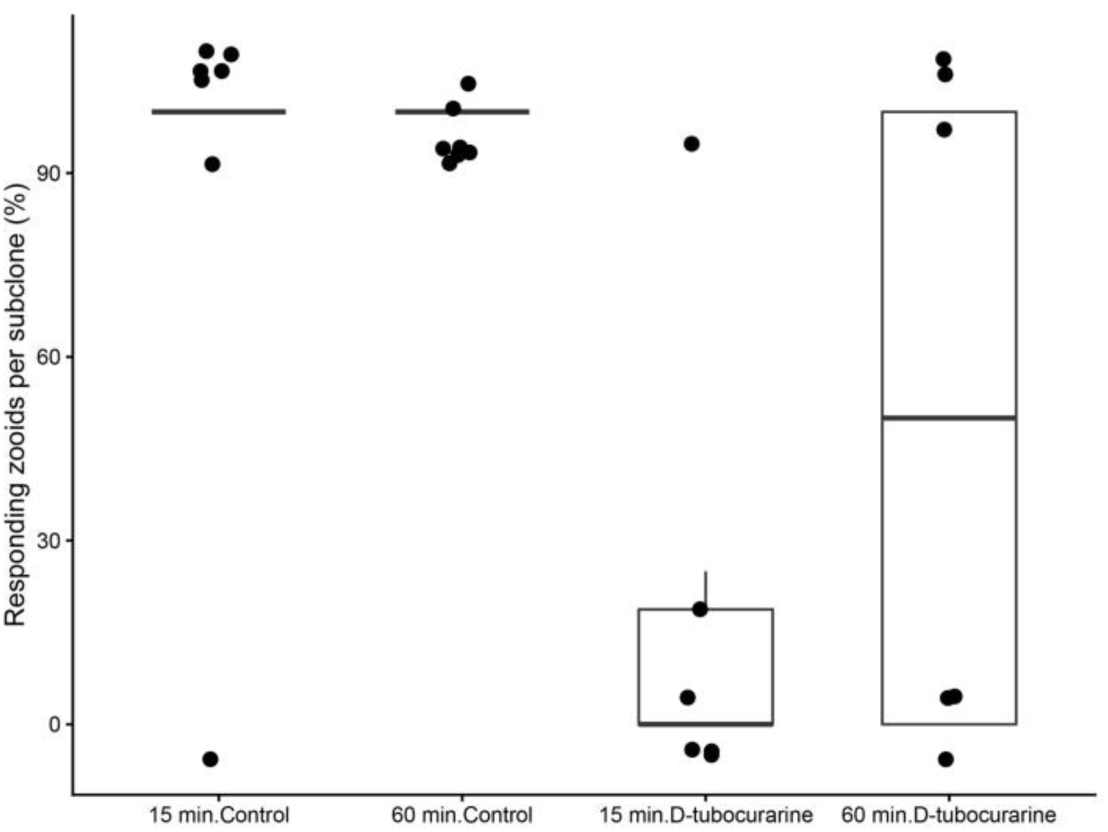
Proportion of responding zooids following focal water-jet stimulation. The fraction of zooids showing an oral siphon contraction in response to focal seawater stimulation was calculated for each control and D-tubocurarine- treated subclone at 15 and 60 min after injection. Each point represents the proportion of responding zooids within one experimental subclone/series. Zooids that did not respond to the applied stimulus were classified as non-responsive. Control subclones received filtered natural seawater, whereas treated subclones received D-tubocurarine.

**Supplementary Table 1**: **Raw data used for the analysis of the focal water-jet stimulation assay.** The table reports individual Stage A zooid measurements from control and D-tubocurarine-treated subclones before injection and at 15 and 60 min after injection. For each measurement, genotype, subclone/experimental-series identity, treatment condition, time point, response threshold, and response outcome are provided. `Threshold_kPa` indicates the pressure required to evoke oral siphon contraction in responding zooids. Measurements without a numerical response threshold were classified as non-responsive (`Responder = 0`) and assigned `No response` in the normalized outcome field; the original qualitative annotation is retained in the `Raw_outcome` column. Multiple zooids were sampled within each subclone at each time point, and individual zooids were not necessarily the same across time points.

**Supplementary Table 2. HCR RNA-FISH probe.** The table lists the gene-specific probe sets used for HCR RNA-FISH analysis.

